# Cell surface proteomics reveal a strategy for targeting myeloma through combinatorial targeting of TNFRSF8 and TMPRSS11E

**DOI:** 10.1101/2023.04.04.535580

**Authors:** Georgina S.F. Anderson, Ieuan G. Walker, James P. Roy, Michael A. Chapman

## Abstract

Chimeric antigen receptor (CAR) T-cell therapy is a highly effective novel treatment in haematological malignancies that has shown promise as a therapeutic option in multiple myeloma. However, widespread adoption of CAR T-cell therapy in myeloma has been hindered by the challenge of unbiased target antigen identification and selection. As activation of CAR T-cells requires minimal antigen on the cell surface, a major risk of toxicity is destruction of healthy tissue expressing the target protein, i.e. on-target, off-tumour toxicity. Indeed, examination of the myeloma surface proteome demonstrated that there was no single target that was completely unique to myeloma cells. One approach to achieve target specificity is to require simultaneous expression of two proteins on the target cells, so-called AND-gate targeting. To identify potential AND-gate combinations for myeloma, we devised an algorithm to prioritise pairings that exhibited pan-myeloma expression and no overlapping expression in vital healthy tissue, as predicted by proteomics. Through this approach, we identified over 600 combinations. To minimise the risk of exhaustion or priming by CAR T-cells, any combination whereby one of the two antigens was expressed in T-cells was also excluded, leading to the prioritisation of 144 candidate pairings. This demonstrates the potential for AND-gating to expand the repertoire of CAR T-cell targets for myeloma. We evaluated one of these candidate pairings, TMPRSS11E and TNFRSF8, *in vitro*. Activation in the Jurkat cell line co-expressing a suboptimal CAR against TNFRSF8 and a chimeric costimulatory receptor (CCR) against TMPRSS11E was markedly enhanced following co-culture with a dual-target positive myeloma cell line compared with single-target positive K562, demonstrating improved discrimination between tumour and non-tumour cells.

## Introduction

Multiple Myeloma (MM) is a haematological malignancy characterised by the clonal expansion of plasma cells within the bone marrow. Despite recent therapeutic advances, including the introduction of monoclonal antibodies, antibody drug conjugates, and new generations of immunomodulatory agents and proteasome inhibitors, MM is still considered to be an incurable disease and nearly all patients will experience relapse. As such, there is an unmet need to find treatment options that provide more consistent and durable remissions.

Chimeric antigen receptor (CAR) T-cells have emerged as a highly effective treatment option in the B cell malignancy setting [1] and as such, have been extensively investigated in the context of MM. CAR T-cells are patient-derived T-cells that have been genetically engineered to express a synthetic immunoreceptor reprogramming them to eliminate the malignant cells. This receptor consists of an extracellular antigen-recognition domain, typically single-chain variable fragments (scFvs) derived from antibodies, fused to intracellular co-stimulatory domains (such as CD28 or 4-1BB) and activation domains (CD3ζ) via a hinge and transmembrane region.

The success of CD19-targeting CAR T-cells has been in part due to the lineage-restricted expression of the target antigen and the translation of this success to other cancers has been hindered by the lack of unique tumour antigens. Proposed targets are invariably also expressed on healthy tissue causing CAR T-cells to attack these non-malignant cells, resulting in on-target, off-tumour toxicities. These can range from the reasonably mild, such as the transient nail loss observed during treatment with a GPRC5D-targeting CAR [2] and B cell aplasia with CD19-targeted CAR T-cell therapies[3], to the much more severe. In a phase 1 dose-escalation study for GPRC5D-targeted CAR T-cell therapy, two patients receiving the highest dose developed grade 3 cerebellar disorder[2]. This was most likely the result of neurological tissue expression of the target antigen, with GPRC5D mRNA detected in the inferior olivary nucleus [2]. Even patients treated with CAR T-cells against BCMA, a purportedly B and plasma-cell exclusive protein [4], have developed on-target, off-tumour toxicities, displaying Parkinson’s-like symptoms attributed to expression in the basal ganglia [5]. These toxicities may even prove fatal, as was the case for a patient with metastatic colon cancer treated with a HER2-targeted CAR product. Within 15 minutes of the infusion, the patient began to experience respiratory distress and after several cardiac arrests, died just five days later. The cause of this toxicity was likely reactivity of the infused CAR T-cells against the low levels of HER-2 in normal lung tissue [6].

One emerging approach to mitigate these on-target, off-tumour toxicities is to use combinatorial antigen recognition, or so-called ‘AND-gates’, whereby the CAR T-cell must recognise two different antigens for target cell elimination. Providing there is no overlapping expression of these antigens, normal tissue is thus spared. There are two main types of AND-gates, relying on either simultaneous or sequential signalling. Split CARs fall into the first category and require the co-expression of a poorly active or suboptimal CAR (containing only the CD3ζ activation domain) targeting one antigen and a chimeric costimulatory receptor specific to a second antigen. Engagement of the target antigen by the CAR alone is insufficient to reach the activation threshold and only through the simultaneous signalling of the chimeric co-stimulatory receptor, is full T-cell activation achieved [7]. Sequential AND-gates rely on a synthetic receptor (SynNotch) to regulate expression of a CAR construct against a second target. The synthetic receptors consist of an scFv fused to part of the Notch receptor and a transcription factor. Engagement of its target antigen results in the release of the transcription factor, which then drives expression of the CAR construct[8].

Logic-gating relaxes the requirement for truly unique tumour antigens. However, identifying target combinations remains a limiting factor in the development of novel CAR T-cell therapies. We have previously reported the characterisation of the MM cell surface proteome [9]. In this study, we integrate this dataset with a normal tissue proteomic dataset [10] to determine a comprehensive list of AND-gate combinations for MM and present TMPRSS11E and TNFRSF8 as an exemplar pairing.

## Methods

### Combinatorial target selection

All data analysis for this study was performed using R (version 4.2.2) [11]. All code used in the generation of this manuscript will be uploaded to the following repository on acceptance in a peer reviewed journal (https://github.com/ieuangw/AND_GATE). Protein expression data for primary MM was obtained from our previous study [9] and for normal tissue from the Human Proteome Map (HPM) [10]. Proteins were annotated according to Uniprot for topological domains and extracellular domain length. Proteins were only included for further analysis if they were identified in the primary MM dataset, contained a transmembrane domain and an extracellular domain length greater than one. A 2-dimensional vector was then created for each possible AND-gate pair and screened against the HPM dataset. A Boolean logic approach was used, whereby combinations were discarded if the vector was detected in any tissue (excluding haematopoietic tissue). To filter for T-cell expression, both proteins in each pairing were annotated according to the HPM dataset as either present or absent in CD4 and CD8 T-cells. Any pairings with T-cell expression, for either target antigen, were excluded.

### CAR constructs

DNA fragments (Thermo Fisher) containing scFvs targeting TNFRSF8 or TMPRESS11E were cloned into pSLCAR-CD19-BBz (Addgene #135992) using NEB HiFi DNA assembly, replacing the FMC63 scFv. The 3XFLAG tag was replaced by Q5^®^ site-directed mutagenesis (NEB) with either a Myc-tag or Strep-tag^®^II. This resulted in a construct that contains as follows: human CD28 signal peptide, an affinity tag, anti-TNFRSF8/TMPRSS11E scFv, human CD8α, human CD28 transmembrane domain, human 4-1BB cytoplasmic domain, human CD247 (CD3ζ) cytoplasmic domain and GFP separated by a 2A self-cleaving peptide. For AND-gating, constructs were modified using Q5^®^ site-directed mutagenesis (NEB) to remove either the 4-1BB costimulatory domain (TNFRSF8-CD3ζ) or the CD3ζ activation domain (TMPRSS11E-41BB).

### CAR T-cell reporter assay

A Jurkat *Nur77*-mCherry reporter line was generated by inserting a T2A-mCherry sequence downstream of the *Nur77* gene by homologous recombination as previously described [12]. CAR constructs were inserted into the Jurkat cell line by lentiviral transduction. Lentivirus was prepared using HEK-293T and added to target cell lines with polybrene (8µg/ml), followed by spinoculation at 750g for 30 minutes. CAR-expressing Jurkat cells were co-cultured with target cells at a 1:1 ratio for 20-24h before flow cytometric analysis of mCherry and CD69 expression. All cells were grown in Roswell Park Medical Institute (RPMI) 1640 medium (Life Technologies) supplemented with 10% fetal bovine serum (FBS; S. American Origin, Cytiva) and 1% GlutaMax (Life Technologies). XG1 cells were supplemented with IL-6 (Roche, 3ng/ml).

### Flow cytometry

Flow cytometry was performed using a LSRFortessa with a high-throughput plate reader. The following reagents were used: anti-MYC-PE (clone 9B11, Cell Signaling Technology), THE™ NWSHPQFEK Tag-biotin (Cambridge Bioscience), anti-CD69-APC (clone FN50, BioLegend), streptavidin-APC (eBioscience), anti-TNFRSF8-PE (clone BY88, BioLegend), anti-TMPRSS11E-PE (clone TM191, BioLegend), DAPI and LIVE/Dead™ Fixable Violet (Thermo Fisher). Data was analysed using FlowJo (Beckton Dickinson, Version 10.8.1). Viable cells were identified by the exclusion of a live/dead stain and selected for downstream analysis following exclusion of doublets and debris.

Jurkat cells expressing the CAR transgenes were identified as GFP+ and mCherry and CD69 expression was measured as the mean fluorescence intensity (MFI). Expression of TNFRSF8 and TMPRSS11E in target cell lines is reported as the ratio of the fluorescence intensity over an isotype-matched control.

## Results

### Identification of combinatorial antigen targets

Antigen expression on the cell surface is essential for therapy. Therefore, to generate our list of combinatorial antigen targets, we first identified cell surface proteins with a targetable extracellular domain. We explored our previously published PMP dataset [9], which contains all cell surface proteins identified in primary MM by mass spectrometry. We selected proteins that were annotated as having a transmembrane domain and an extracellular domain length of greater than one amino acid, a total of 777 proteins, and determined there were a total of 287,890 possible combinations for AND-gating on primary MM cells.

From these possible combinations, we filtered for off-tumour expression. The human proteome map provides a comprehensive list of protein expression across multiple histologically normal human cell and tissue types as determined by mass spectrometry [10]. By integrating this dataset with ours, we were able to evaluate antigen combinations by their expression in healthy tissue. Only combinations which had non-overlapping expression in vital tissue (frontal cortex, spinal cord, retina, heart, liver, lung, adrenal gland, gallbladder, pancreas, kidney, oesophagus, colon, rectum, and urinary tract) were retained, reducing our total pairings to 664. We then further excluded any combinations where either target was expressed in T-cells. This was done to address two concerns. Firstly, that CAR T-cells may be ‘primed’ by themselves, enabling the subsequent killing of nearby single-antigen positive cells. And secondly, that the persistent signalling through one of the two constructs may drive CAR T-cells towards a state of exhaustion, greatly diminishing anti-tumour efficacy [13]. This reduced our list from 664 to just 144 combinations. Owing to their small extracellular domains, highly conserved sequences, and complex structures, multi-pass proteins are typically challenging for antibody generation and so we also excluded pairings containing these types of proteins. This left us with a total of 92 AND-gate combinations which were predicted to be highly expressed in MM but were not co-expressed in normal, non-haematopoietic tissue. An overview of our prioritisation strategy is shown (**Fig. 1**).

**Figure 1.**
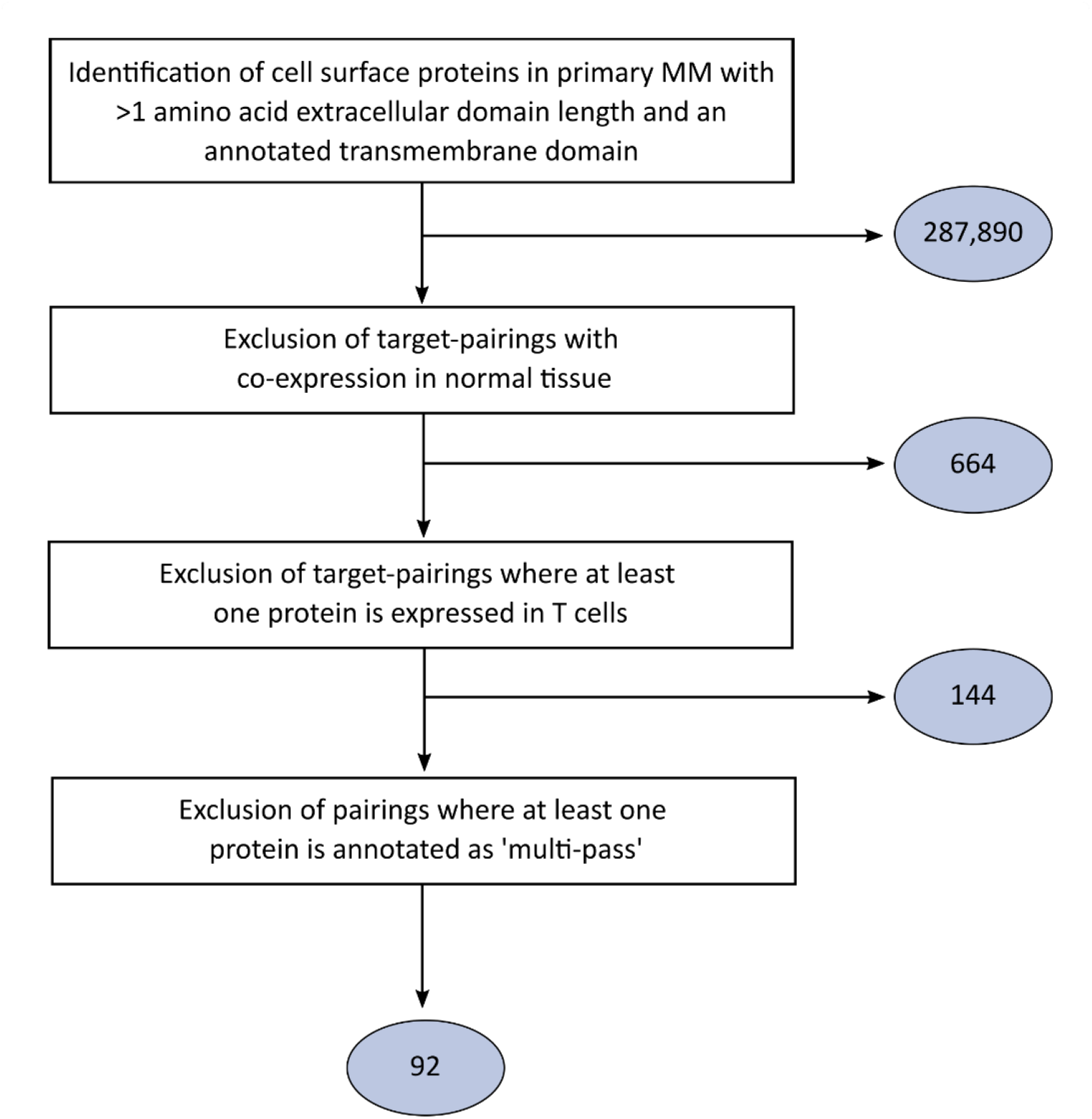
Overview of the algorithm applied to identify multiple myeloma AND-gate pairings. A flow diagram showing the computational pipeline used in this study. The number of pairings remaining after each step are highlighted in the blue ovals.

### The majority of AND-gate pairings are comprised of a small number of proteins

To understand whether specific proteins have multiple partners on MM or rather that the distribution of partners is random, we analysed the number of partner proteins each MM cell surface protein made. After excluding for combinations with overlapping expression in healthy tissue, we observed that only a few proteins formed multiple combinations (**Fig. 2**) and of our top 92 AND-gate pairs, all were comprised of one of four proteins (ITGA2, ITGA3, ITGAV and TMPRSS11E). All four of these proteins have already been described in other cancers as potential biomarkers or therapeutic targets [14-17] and so may represent antigens that are involved in tumorigenesis that would be un-targetable using traditional CAR T-cell therapy due to low expression in normal tissue. TMPRSS11E was the most common partner (66/92), most likely a result of its low predicted expression across multiple tissue types which allows a greater flexibility in antigen-pairing.

**Figure 2.**
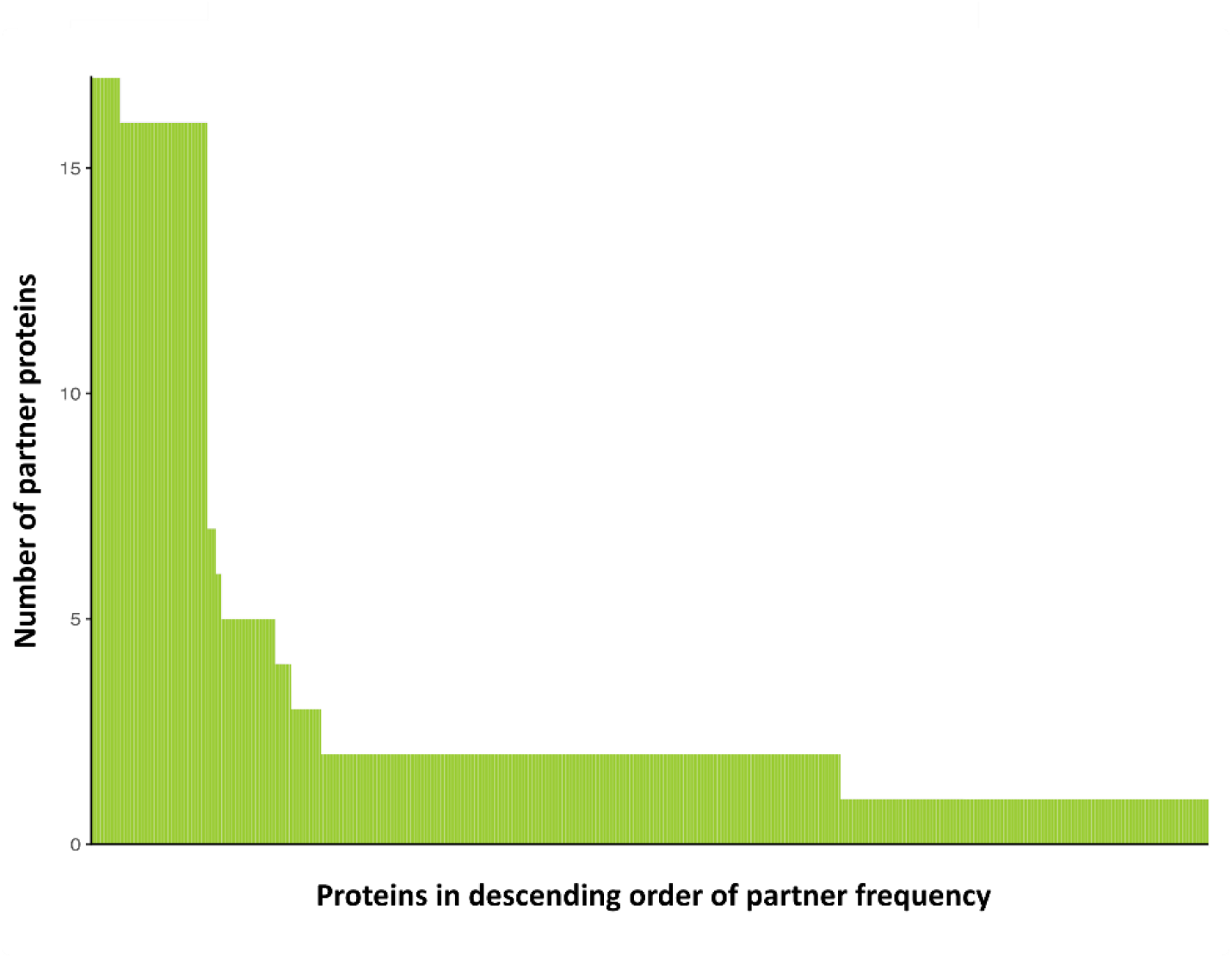
Only a few proteins form repeated target pairings. The frequency of partner proteins in our AND-gate pairings.

### Prioritisation of AND-gate combinations

We wanted to prioritise target combinations from amongst our 92 potential pairs. Various options exist to do this. For instance, to mitigate the risk of antigen escape, results from genome wide CRISPR/Cas9 screens could be incorporated to predict target essentiality [18]. To improve the likelihood of raising scFvs against candidate antigens, one might prefer to prioritise proteins that are predicted to be highly immunogenic. Alternatively, target candidates that are associated with therapeutic resistance or known to be upregulated in relapsed/refractory MM may be more reflective of the tumours of patients undergoing CAR T-cell therapy. To identify an exemplar antigen-pair, we prioritised candidates that had not been previously described as CAR T-cell targets for MM, had commercially available antibodies available for target validation and scFv sequences available.

From these candidates, we chose TNFRSF8 (CD30) and TMPRSS11E for further evaluation. Both TMPRSS11E and TNFRSF8 were expressed in all eight primary samples by PMP and at comparable levels to two other MM CAR T-cell targets (**Fig. 3A**). As predicted by our algorithm, there was no overlapping expression in normal tissue (**Fig. 3B**).

**Figure 3.**
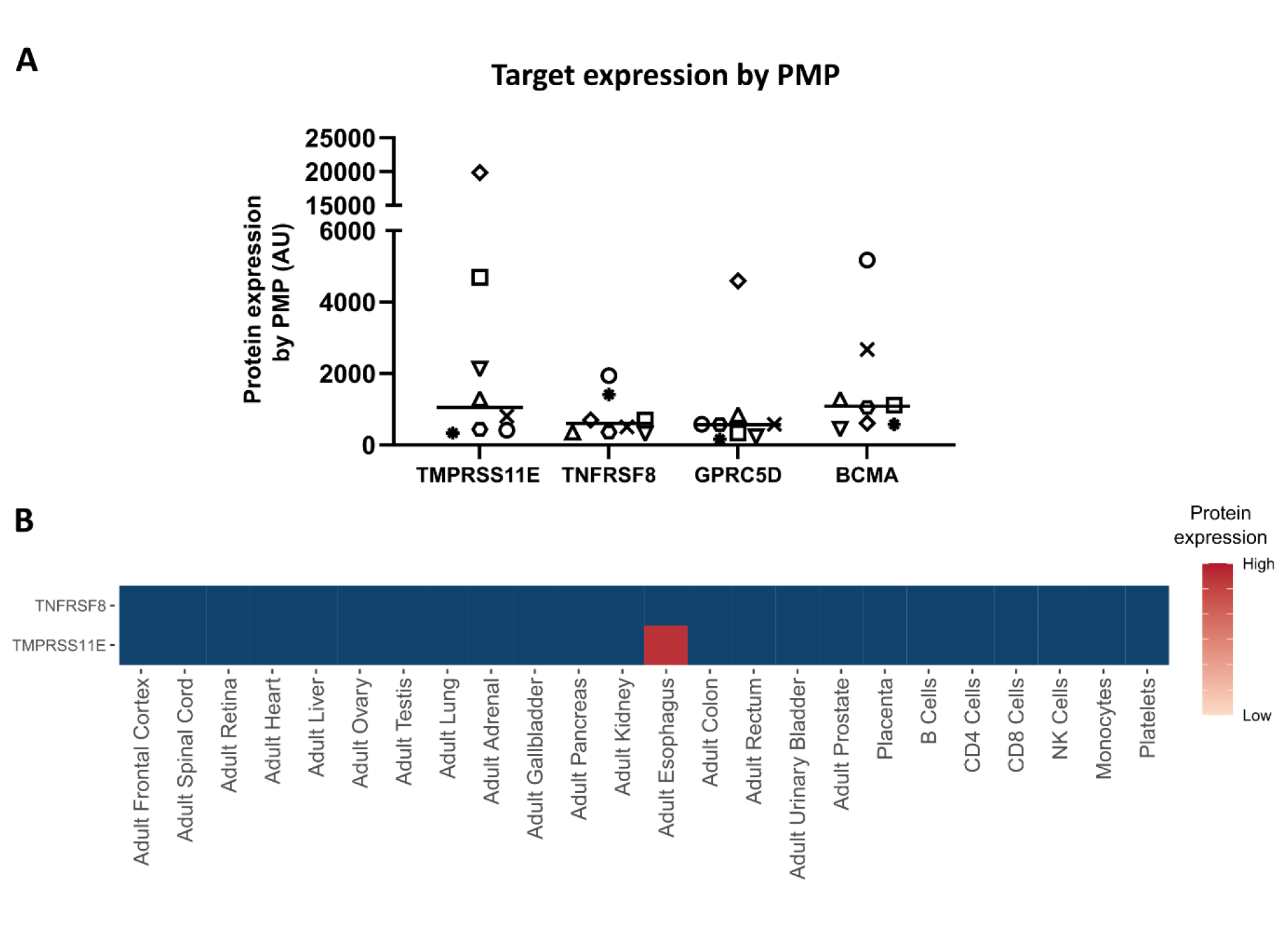
TMPRSS11E and TNFRSF8 form a viable AND-gate combination. (A) TNFRSF8 and TMPRSS11E expression by plasma membrane profiling (PMP) in primary multiple myeloma cells (n=8). Each unique symbol identifies a patient. Included for comparison are two well-described candidate targets for anti-MM CAR T therapy, GPRC5D and BCMA. (B) Heatmap showing expression of TMPRSS11E and TNFRSF8 in normal healthy tissues by whole cell proteomic profiling (Human Proteome Map [10]). Blue indicates not detected.

### Optimal Jurkat activation requires engagement of both the CAR and chimeric co-stimulatory receptor

A major disadvantage with the SynNotch system is that the CAR construct is not immediately degraded following the loss of signalling through the primary synthetic receptor. As a consequence, there is still a risk of cytolytic activity towards neighbouring single-antigen positive cells [8]. For this reason, we chose to proceed with a split-CAR design and tested these constructs in a Jurkat cell line model of T-cell activation.

Our anti-TNFRSF8 CAR construct (TNFRSF8-CD3ζ) was comprised of a CD28 signal peptide, followed by an anti-TNFRSF8 scFv, CD8α hinge, CD28 transmembrane domain and a CD3ζ activation domain. The costimulatory anti-TMPRSS11E receptor (TMPRSS11E-41BB) was similarly designed, but the intracellular portion contained a 4-1BB costimulatory domain as opposed to the CD3ζ signalling domain (**Fig 4A**). CAR constructs were stably transduced into a fluorescent activation reporter cell line (*Nur77*-mCherry Jurkat). In this cell line, T2A-mCherry has been inserted downstream of *Nur77*, an immediate early-response gene that is upregulated following anti-CD3 stimulation. As fluorescent protein expression in *Nur77* reporter cells has been shown to be indicative of signal strength [19], this provides a simple and quantitative readout of CD3-dependent activation in a T cell model.

**Figure 4.**
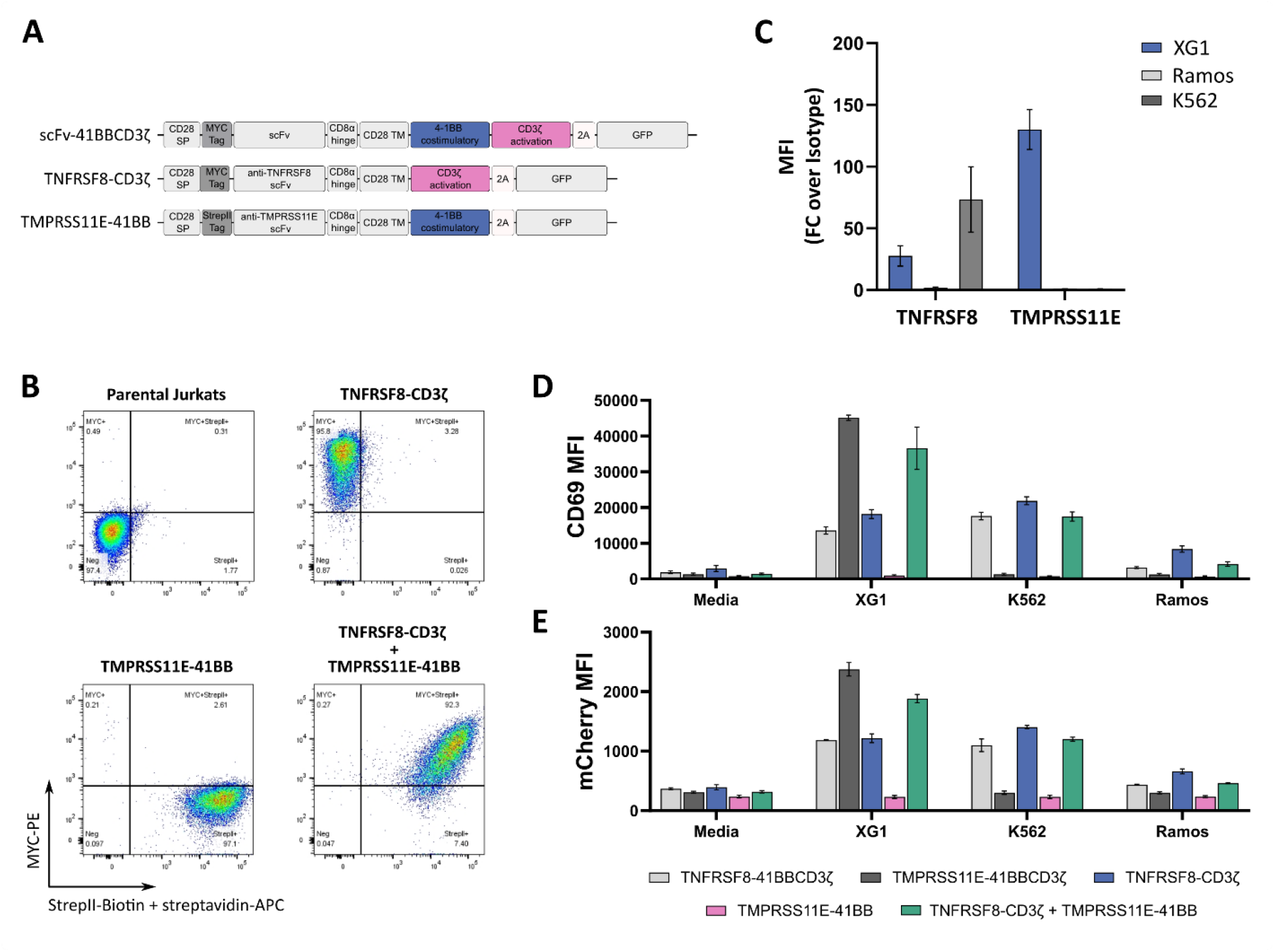
Jurkat reactivity against dual-target positive cells is enhanced by the co-expression of a chimeric co-stimulatory receptor. A. Schemata of the different CAR constructs used in this study. TNFRSF8/TMPRSS11E-41BBCD3ζ: A CD28 signal peptide (SP) is followed by the scFv, CD8α hinge region, CD28 transmembrane domain (TM), 4-1BB costimulatory domain and a CD3ζ activation domain. TNFRSF8-CD3ζ: suboptimal car lacking the 4-1BB co-stimulatory domain. TMPRSS11E-41BB: chimeric co-stimulatory receptor lacking the CD3ζ activation domain. B. Expression of TNFRSF8 and TMPRSS11E by flow cytometry in XG1, Ramos and K562. Expression was calculated as the median fluorescence intensity of the antibody relative to the isotype control (FC = fold change, n≥3). C. Cell surface expression of CAR constructs in Jurkat-cells by flow cytometry. TNFRSF8-CD3ζ+ cells were detected using a PE-conjugated anti-MYC antibody and TMPRSS11E-41BB+ cells using a biotinylated-anti-Strep-tag™II antibody and streptavidin-APC. D. CAR-expressing Jurkats were co-cultured at a 1:1 ratio with the following target cells: TMPRSS11E+TNFRSF8+ XG1, TMPRSS11E-TNFRSF8+ K562 or TMPRSS11E-TNFRSF8- Ramos for 20-24h. Jurkat activation was measured by the mean fluorescence intensity (MFI) of CD69 or mCherry (inserted in-frame with *Nur77*) in CAR+ cells, n=2. Error bars are ± SD of biological replicates.

Second generation CAR constructs, which contain both the costimulatory and activation domains, were included as positive controls. Cell surface expression of the constructs was confirmed by flow cytometry (**Fig. 4B**) using antibodies against the Myc-tag or Strep-tag^®^II inserted in-frame between the CD28 signal peptide and the scFv. As shown, expression of the CAR and chimeric co-stimulatory receptor was comparable between single- and dual-targeted Jurkats.

To evaluate the activity of these constructs, CAR-expressing Jurkats were co-cultured *in vitro* with the MM cell line XG1 (TMPRSS11E+TNFRSF8+) or the erythroleukemic line K562 (TMPRSS11E-TNFRSF8+). Ramos, a B-cell lymphoma line, which expresses neither antigen, served as a negative control (**Fig 4C**). Jurkat activation was measured by both mCherry (Nur77) and CD69 upregulation. CD69 was included as, unlike Nur77, it may be upregulated by TCR-independent mechanisms and so enables us to identify any activation independent of the CD3ζ domain [20].

We first validated that the scFvs were capable of inducing Jurkat activation when incorporated into second generation CAR constructs. As anticipated, TNFRSF8-41BBCD3ζ expressing Jurkats upregulated both CD69 and mCherry (**Fig. 4D and E**) following co-culture with XG1 and K562. TMPRSS11E-41BBCD3ζ+ Jurkats, however, were only activated by XG1. Co-culture with the target-negative Ramos failed to induce activation in Jurkats expressing either construct.

We next sought to determine whether we could reduce Jurkat activation against single-target cells using our split-CAR system. As anticipated, removal of the CD3ζ activation domain in the TMPRSS11E-targeted chimeric co-stimulatory receptor completely abrogated the ability of the construct to activate Jurkats (**Fig 4D and E**). Even when expressed by itself, signalling through the suboptimal TNFRSF8-CAR was not reduced following removal of the 4-1BB domain and we observed modest activation following co-culture with either XG1 or K562, as determined by CD69 expression (MFI of 18711 ±1256(AU) for XG1 vs 21891 ±1097(AU) for K562, mean ±SD, n=2). However, co-expression of both the CAR and chimeric co-stimulatory receptor markedly enhanced activation, with a two-fold increase in CD69 cell surface expression (18,171 ±1256(AU) vs 36,631 ±5916(AU) for TNFRSF8-CD3ζ alone vs TNFRSF8-CD3ζ+TMPRSS11E-41BB against XG1, mean ±SD, n=2). This robust enhancement was only observed following co-culture with the dual-target positive XG1, and not with the single-target positive K562 (21891 ±1097(AU) vs 17,487 ±1278(AU) for TNFRSF8-CD3ζ alone vs TNFRSF8-CD3ζ+TMPRSS11E-41BB, mean ±SD), demonstrating that engagement of both antigens was required for the augmented activation.

## Discussion

The development of CAR T-cells for non-B cell malignancies has been hindered by the lack of truly unique tumour cell surface proteins, with severe on-target, off-tumour toxicities a major risk [2, 5, 6]. By improving malignant versus healthy tissue discrimination, combinatorial antigen targeting mitigates these toxicities. In this study, we present a computational approach to identifying antigen-pairs that are co-expressed only in MM and not vital healthy tissue or T cells; and report the finding of 144 such potential combinations. Although this is unlikely to be an exhaustive list, this represents a significant contribution to our potential armoury against MM.

From this list, we identified TMPRSS11E and TNFRSF8 as a candidate pairing for AND-gate CAR T-cell therapy. Whilst TMPRSS11E has not been previously described in MM, TNFRSF8 has. However, owing to low expression on patient samples by flow cytometry, it has been previously considered unattractive as a single-target antibody therapy [21]. However, CAR T-cells can recognise and eliminate target cells expressing extremely low antigen levels, far below that which is detectable by conventional flow cytometry [22]. As TNFRSF8 expression was detectable by mass spectrometry across all eight primary MM samples at comparable levels to the CAR T-cell target GPRC5D (**Fig 2A**), we anticipate that expression would be sufficient for CAR T-cell mediated killing.

By splitting the activation and costimulatory domain across two receptors, full T-cell activation relies on the simultaneous engagement of TNFRSF8 and TMPRSS11E. We show that Jurkats expressing the CAR construct alone had significantly reduced activation compared with cells co-expressing both the CAR and chimeric co-stimulatory receptor, and that optimal activation was dependent on the engagement of both target antigens. Although we were unable to abrogate signalling through the suboptimal CAR, activation *in vitro* by single-target cells is unlikely to be indicative of cytolytic capability *in vivo*. First generation CAR T-cells also only contained a CD3ζ cytoplasmic domain, and despite demonstrating efficacy *in vitro*, had limited anti-tumour activity *in vivo* [23]. Similarly, in a study by Lanitis *et al*, T-cells expressing suboptimal CARs were activated by and demonstrated cytolytic activity against single-target expressing cell lines *in vitro* [24]. However, *in vivo*, these T-cells exhibited significantly reduced activity against both single-antigen and dual-antigen positive tumour cells compared to T-cells co-expressing the chimeric co-stimulatory receptor. Importantly, the authors also demonstrated that the enhanced specificity of these split-CARs was not at the cost of potency. For future studies, it may be prudent to include additional measures of function, such as cytokine release and proliferative capacity, to truly validate split-CARs *in vitro* [24, 25].

It is important to note that the split-CAR constructs used in this study have not been optimised, and as such there is potential to further improve specificity for dual-target cells. By reducing the affinity of the scFv used in the CAR construct, Kloss *et al*, were able to completely abolish reactivity against single-target cells [7]. Other modifications could include substituting the CD28 transmembrane domain with CD8α, preventing signal amplification through dimerization [26] and by incorporating the CD28 co-stimulatory domain alongside 4-1BB in the chimeric co-stimulatory receptor to further enhance signalling [27].

Although steps may be taken to minimise signalling through the CAR, there remains a modest risk for activity against single-antigen positive cells and so antigen assignment to the CAR or chimeric co-stimulatory receptor is crucial. Although TNFRSF8 was predicted to have no expression across all tissue types in the HPM, it is well-known to be expressed in a subset of activated T and B cells [28].

Reactivity against these cells would be much more tolerable than against oesophageal tissue (**Fig 2B**), and so we reasoned that this protein should be assigned as the CAR antigen. As TNFRSF8 expression is upregulated following CD3/CD28 stimulation, we may observe some degree of fratricide during the CAR T-cell manufacturing process, but we anticipate that this may not be detrimental to CAR T-cell efficacy. In fact, due to its regulatory role, depletion of TNFRSF8+ cells may enhance cytolytic activity [29] and minimise the risk of priming against single-antigen positive cells.

That TNFRSF8 was not flagged as being expressed in T-cells by the HPM dataset is an important caveat of this study. The HPM relies on whole cell proteomics, which may under-estimate the expression of plasma membrane proteins [30]. Furthermore, proteins that are transiently upregulated or are only expressed in subset of cells within a tissue, such as the case for TNFRSF8, may not be captured by this type of analysis. By integrating additional normal cell proteomic datasets, such as the human protein atlas [31], we may be able to improve our confidence in scoring off-tumour expression. Whilst we anticipate that TNFRSF8 expression on a subset of T and B cells represent an acceptable safety risk, other false negatives may be much more deleterious. Further improvements could also be made by incorporating single-cell transcriptomic datasets. It is imperative in split-CAR targeting that both antigens are co-expressed across all tumour cells. As MM shows great intra-patient heterogeneity, it would be highly advantageous to exclude any combinations not ubiquitously expressed across subclones.

In summary, by applying a computational approach, we have identified several potential AND-gate pairings and evaluated one such combination in a Jurkat model of activation. Although additional studies are required to fully validate this combination, the current study presents a proof-of-concept for a pipeline to discover AND-gate CAR T-cell targets for MM.

